# *Ms4a4a* loss reprograms amyloid-associated microglia and limits dense-core plaque-associated tau spreading

**DOI:** 10.64898/2026.07.22.740167

**Authors:** Emma P. Danhash, Shao-Yu Fang, Jacob A. Marsh, Ricardo D’Oliveira Albanus, Anthony C. Verbeck, Guangming Huang, Shih-Feng You, Erin E. Franklin, Richard Perrin, Wade K. Self, David M. Holtzman, Celeste M. Karch

**Author notes:** Corresponding Author: Celeste M. Karch, Department of Psychiatry, Washington University in St Louis, 660 South Euclid Ave., MSC 8134-0096-06, St. Louis, MO 63110-1010.

## Abstract

**INTRODUCTION:** Microglia regulate amyloid plaque-associated microenvironments that contribute to downstream tau pathology in Alzheimer’s disease (AD). Variants within the *MS4A* locus are strongly associated with AD risk and resilience and are linked to microglial biology; however, the functional role of *MS4A4A* in plaque-associated tau pathology remains poorly understood.

**METHODS:** Single-nucleus RNA sequencing (snRNA-seq) was performed on hippocampi from non-transgenic, *Ms4a4a* knockout (4A-KO), 5xFAD, and 5xFAD 4A-KO mice at 6 months of age. To assess plaque-associated tau pathology, AD-derived tau aggregates were injected into the hippocampus of 5xFAD and 5xFAD 4A-KO mice at 6 months, and histological analyses were performed 3 months later.

**RESULTS:** Amyloid pathology was the dominant driver of microglial state transitions, while *Ms4a4a* loss selectively remodeled activated microglial transcriptional programs enriched for interferon, lysosomal, autophagic, and proteostatic pathways. Activated microglia from 5xFAD 4A-KO mice exhibited altered expression of genes linked to immune signaling and protein handling. Following AD-tau inoculation, *Ms4a4a* loss did not significantly alter overall phospho-tau burden but selectively reduced dense-core plaque-associated neuritic plaque tau (NP-tau), particularly in the contralateral hemisphere. This phenotype was strongest surrounding X-34-positive fibrillar plaques and occurred without major changes in plaque-associated microgliosis.

**DISCUSSION:** These findings identify *Ms4a4a* as a regulator of plaque-associated microglial programs linked to NP-tau accumulation in the amyloid-bearing brain. More broadly, this work supports a model in which AD resilience-associated microglial pathways selectively shape plaque-associated microenvironments that promote downstream tau pathology.

## Background

Alzheimer’s disease (AD) is characterized by the accumulation of extracellular amyloid beta (Aβ) plaques and intracellular neurofibrillary tangles (NFT) composed of hyperphosphorylated tau aggregates. While amyloid deposition is thought to initiate disease pathogenesis, tau pathology is more closely associated with neurodegeneration and cognitive decline (van der Kant, Goldstein et al. 2020). Increasing evidence suggests that amyloid plaques create local microenvironments that promote neuritic injury, inflammatory activation, and the accumulation of phosphorylated tau within plaque-associated dystrophic neurites, termed neuritic plaque tau (NP-tau) (Dickson and Vickers 2001, Meyer-Luehmann, Spires-Jones et al. 2008, Condello, Yuan et al. 2015, Sadleir, Kandalepas et al. 2016, He, Guo et al. 2018). However, the mechanisms linking amyloid deposition to local tau pathology remain incompletely understood. Among the cellular populations surrounding amyloid plaques, microglia have emerged as critical regulators of plaque formation, compaction, proteostatic responses, and downstream neurodegenerative processes (Baik, Kang et al. 2016, Hansen, Hanson et al. 2018, Spangenberg, Severson et al. 2019, Casali, MacPherson et al. 2020, Choi, Wang et al. 2023, Baligacs, Albertini et al. 2024). Genetic studies further implicate microglial pathways as major contributors to AD risk and resilience (Treusch, Hamamichi et al. 2011, Guerreiro, Wojtas et al. 2013, Jonsson, Stefansson et al. 2013, Karch and Goate 2015, Efthymiou and Goate 2017, Sims, van der Lee et al. 2017, Nott, Holtman et al. 2019, Novikova, Kapoor et al. 2021, Bellenguez, Kucukali et al. 2022, Brase, You et al. 2023, Sun, Victor et al. 2023), supporting a model in which microglial response to amyloid pathology shapes the transition from Aβ accumulation to tau-mediated neurodegeneration.

NP-tau accumulates within dystrophic neurites surrounding amyloid plaques and is particularly enriched near dense-core fibrillar plaques (He, Guo et al. 2018), suggesting that local plaque-associated cellular responses influence tau pathology. Microglia tightly cluster around plaques where they regulate plaque compaction, lysosomal pathways, and proteostatic responses that may influence neuronal injury and pathogenic protein accumulation (Condello, Yuan et al. 2015, He, Guo et al. 2018, Leyns, Gratuze et al. 2019, Sadleir, Gomez et al. 2025). Disease-associated microglial states enriched near plaques exhibit transcriptional programs linked to interferon signaling, lysosomal activation, and cellular stress responses (Sun, Victor et al. 2023, Avey, Ng et al. 2025). Targeting microglia via pharmacological ablation, genetic knockout, or introduction of AD-associated variants alters NP-tau seeding and/or spreading (Leyns, Gratuze et al. 2019, Delizannis, Nonneman et al. 2021, Gratuze, Chen et al. 2021, Wang, Fan et al. 2022, Chen, Song et al. 2024). However, the specific microglial pathways that regulate plaque-associated tau accumulation remain poorly understood.

Genome-wide association studies have identified the *MS4A* locus as a major regulator of AD risk and resilience (Novikova, Kapoor et al. 2021, Wightman, Jansen et al. 2021, Bellenguez, Kucukali et al. 2022). Among genes within this locus, *MS4A4A* is highly enriched in microglia and strongly associated with cerebrospinal fluid soluble TREM2 levels (Deming, Filipello et al. 2019, Brase, You et al. 2023, You, Brase et al. 2023, Wang, Nykanen et al. 2024), implicating MS4A4A in microglial activation and innate immune signaling. We previously demonstrated that loss of *Ms4a4a* in 5xFAD mice reduces amyloid plaque burden, alters plaque compaction, and modifies plaque-associated microglial responses at 6 months of age (Danhash, Verbeck et al. 2025), supporting a role for *Ms4a4a* in regulating microglial responses to amyloid pathology. More recently, emerging studies suggest that MS4A family proteins modulate microglial effector programs linked to lysosomal function, inflammatory signaling, and proteostatic pathways (Rosner, Sun et al. 2026). However, whether *Ms4a4a*-dependent microglial programs influence plaque-associated tau pathology remains unknown.

A major unresolved question in AD is how microglial states associated with amyloid pathology influence the local plaque microenvironment to promote downstream tau accumulation and neurodegeneration. In particular, it remains unclear whether AD resilience-associated microglial pathways regulate NP-tau accumulation independently of broad reductions in amyloid burden. Defining how microglial programs shape plaque-associated tau pathology may provide insight into mechanisms linking amyloid deposition to downstream neurodegenerative processes and identify therapeutic pathways that selectively modulate pathogenic plaque microenvironments.

In the present study, we combined single-nucleus RNA sequencing, AD-derived tau (AD-tau) inoculation, and histological analyses to define how *Ms4a4a* loss alters microglial states and plaque-associated tau pathology in the amyloid-bearing brain. We demonstrated that *Ms4a4a* loss remodels activated microglial transcriptional programs enriched for interferon, lysosomal, and proteostatic pathways. In parallel, *Ms4a4a* loss selectively reduces dense-core plaque-associated NP-tau accumulation following AD-tau inoculation without substantially altering overall phospho-tau burden or late-stage amyloid plaque burden. Together, these findings support a model in which *Ms4a4a*-dependent microglial programs regulate plaque-associated microenvironments linked to NP-tau accumulation during amyloid pathology.

## Methods

### Animals

Animal care and surgical procedures were approved by the Animal Studies Committee of Washington University School of Medicine in accordance with guidelines from the United States National Institutes of Health. 5xFAD (B6.Cg-Tg (APPSwFlLon, PSEN1*M146L*L286V) 6799Vas/Mmjax, Stock number 034848-JAX, MMRRC, The Jackson Laboratory) hemizygous mice were crossed with ^+/-^ mice (generously provided by Alector; constitutive knockout generated using CRISPR/Cas9 to excise exons 1-7 in C57BL/6 mice) to generate 5xFAD x *Ms4a4a*^+/-^ mice (Danhash, Verbeck et al. 2025). Next, 5xFAD x *Ms4a4a*^+/-^ were crossed with *Ms4a4a*^+/-^ mice to obtain 5xFAD x *Ms4a4a*^-/-^ (5xFAD 4A-KO) mice and 5xFAD x *Ms4a4a*^+/+^ (5xFAD 4A-WT) littermates. Genotype was confirmed using primers to determine the presence of *Ms4a4a* and the 5xFAD transgene (**Supplemental Table 1**).

### Mouse hippocampus nuclei isolation, library preparation, and single nucleus RNA sequencing (snRNAseq)

Hippocampi were dissected from 6-month-old male and female mice and flash-frozen on dry ice. Samples included non-transgenic *Ms4a4a* WT (n=2), non-transgenic *Ms4a4a* KO (n=2), 5xFAD-*Ms4a4a* WT (n=6; 3 females), and 5xFAD-*Ms4a4a* KO (n=6; 3 females). Nuclei were isolated as previously described with some modifications (Brase, You et al. 2023). Briefly, frozen hippocampi were homogenized in supplemented homogenization buffer (SHB; Homogenization Buffer, 10% spermine, 10% spermidine; 1% RNASIN and 0.1 mM EDTA), and nuclei were purified by density gradient centrifugation using iodixanol solutions of varying concentrations. Isolated nuclei were visually inspected by trypan blue staining to assess nuclear integrity and cell lysis. Following counting, nuclei were diluted to 1000 nuclei per μl for snRNA-seq. SnRNA-seq libraries were generated from mouse hippocampal nuclei using the 10x Genomics Chromium platform. Approximately 10,000 nuclei per sample were loaded onto Chromium single-cell chips, and single-nucleus cDNA libraries were prepared and dual-indexed using the Chromium Single Cell 3⍰ Reagent Kit v3.1 according to the manufacturer’s instructions. Libraries were pooled at equimolar ratios and sequenced on an Illumina NextSeq 2000 platform by the McDonnell Genome Institute at Washington University in St. Louis. Libraries were sequenced to a saturation of >30%.

### snRNAseq data analysis and downstream analyses on microglial subclusters

Raw count matrices were generated per sample using CellRanger-arc-2.0.2. Initial quality control was performed individually on each of the 16 samples using the Seurat 5.3.1 package in R (v4.2.1). Mitochondrial and ribosomal gene content were calculated per nucleus. Adaptive per-sample thresholds were applied to filter nuclei based on UMI counts, feature counts, and mitochondrial fraction. Of 155,598 total nuclei detected, 132,618 (85.2%) passed the initial QC. Doublet detection was performed using DoubletFinder 2.0.6. Doublet rates ranged from 4-7% across samples, and 7,459 nuclei containing doublets were filtered out, yielding 125,159 singlet nuclei. Ambient RNA contamination was subsequently estimated and corrected on a per-sample basis using DecontX.

Data integration, normalization, variable feature identification, scaling, and principal component analysis (PCA) were performed using the standard Seurat v5.3.1 workflow. The 16 samples were then merged, and batch effects across samples were corrected using Harmony 1.2.0. UMAP dimensionality reduction was conducted using the top 20 PCs, selected based on inspection of an elbow plot. Clustering was performed using the FindNeighbors and FindClusters functions in Seurat, which employ a k-nearest neighbors (KNN) graph model and the Louvain algorithm, at a resolution of 0.3. Cell types were annotated by integrating Azimuth mouse cortex reference predictions with canonical marker gene expression, and cluster identities were manually assigned. After cell type annotation, the dataset contained nuclei with 73,582 glutamatergic neurons, 8,444 GABAergic neurons, 11,404 microglia, 7,108 astrocytes, 17,841 oligodendrocytes, 4,224 oligodendrocyte precursors cells (OPCs), 447 vascular leptomeningeal cells (VLMCs), and 2,109 other cells. Microglia were then subsetted from the integrated dataset and re-integrated using Harmony. Microglial subclustering was performed using the first 25 PCs at a resolution of 0.3, identifying nine transcriptionally distinct microglial subclusters.

Compositional analysis was performed using propeller, a published method for testing compositional changes in single-cell datasets (Phipson, Sim et al. 2022). Statistical significance was assessed using the ANOVA framework implemented in propeller. Differential gene expression analysis comparing 5xFAD 4A-KO versus 5xFAD 4A-WT mice was performed within each microglial subcluster using the Model-based Analysis of Single-cell Transcriptomics (MAST) framework to assess gene-level statistical significance based on a linear model (Finak, McDavid et al. 2015). Pathway enrichment analysis was conducted separately on upregulated and downregulated differentially expressed genes (p<0.05) within each subcluster using the clusterProfiler package. Gene Ontology (Molecular Function, Cellular Component, and Biological Process), KEGG, and Reactome databases were queried, and enriched pathways were reported at a Benjamini-Hochberg adjusted p<0.05.

### AD-tau aggregate isolation from human brain tissue

AD-tau was isolated from ∼100g of frozen frontal cortex tissue from Washington University Knight ADRC participant brain donors with severe AD dementia (Clinical Dementia Rating® 3) and neuropathology-confirmed high AD neuropathologic change (Thal Aβ phase 5, Braak stage VI, CERAD neuritic plaque score 3) without evidence of synucleinopathy or TDP-43 proteinopathy as previously described (Guo, Narasimhan et al. 2016, Gratuze, Chen et al. 2021). Briefly, the frontal cortical gray matter was dissected and homogenized in a Dounce homogenizer with nine volumes (v/w) of high-salt buffer (10mM Tris-HCl, pH 7.4, 0.8M NaCl, 1mM EDTA, and 2mM dithiothreitol, 1mM PMSF, 1.5μM Pepstatin, 2μM Leupeptin, 3μM TPCK, 3μM TLCK, 1μg/mL Kunitz soybean trypsin inhibitor, 2mM Imidazole, 1mM NaF, 1mM NaOrthovanadate) with 0.1% sarkosyl and 10% sucrose. The homogenate was centrifuged at 10,000×g for 10 minutes at 4°C, and supernatant was collected. Pellets were re-extracted two more times by repeating this process. The three supernatant solutions were filtered, pooled, and stirred for 1.5 hours at room temperature after increasing the sarkosyl concentration to 1%. The sample was then centrifuged at 300,000×g for 1 hour at 4°C. The resulting 1% sarkosyl-insoluble pellet containing pathological tau was washed once then resuspended in PBS (100mL/g gray matter) by passing sequentially through a 27G needle. The suspension was sonicated with 20 pulses for 0.5 second per pulse then centrifuged at 100,000×g for 30 minutes at 4°C. The pellet was resuspended in PBS at one-fifth to one-half of the pre-centrifugation volume, sonicated with 20-60 pulses for 0.5 second per pulse, then centrifuged at 10,000×g for 30 minutes at 4°C. The final supernatant containing AD-tau was frozen at -80°C until use.

### Stereotactic intracerebral injections of AD-tau

Stock AD-tau was thawed on ice then diluted to a concentration of 0.4μg/μL in sterile PBS. AD-tau was sonicated in a water bath sonicator (QSonica, Q700) for 30 seconds at 60% amplitude at 4°C, then placed on ice before injections. Six-month-old female 5xFAD 4A-KO and 5xFAD 4A-WT mice were anesthetized with isoflurane and immobilized using a stereotactic frame (David Kopf Instruments). Using a Hamilton syringe (Hamilton; syringe 80265–1702RNR; needle 7803–072), 2μg AD-tau (1μg per injection site) was unilaterally injected into the dentate gyrus (bregma: 2.5mm; lateral: 2.0mm; depth: 2.2mm) and overlying cortex (bregma: 2.5mm; lateral: 2.0 mm; depth: 1.0 mm). Mice were allowed to recover on a heating pad at 37°C and monitored for 72 hours after surgery.

### Mouse brain tissue preparation

Three months post-AD-tau injection, mice were anesthetized with sodium pentobarbital and perfused with ice-cold PBS. Whole brains were extracted and fixed in 4% paraformaldehyde (w/v) for 48 hours followed by 30% sucrose in PBS at 4°C. Coronal sections (40μm) were cut on a freezing-sliding microtome, and slices were collected and stored in cryoprotectant solution (0.2 M phosphate-buffered saline, 30% sucrose, and 30% ethylene glycol) at -20°C.

### 3,3’-diaminobenzidine (DAB) staining for NP-tau and amyloid plaque quantification

Brain sections from each animal were incubated in 0.3% H_2_O_2_ in Tris-buffered saline (TBS) for 10 minutes then blocked in 3% milk in TBS containing 0.25% Triton-X100 (TBS-X) for 30 minutes. Primary antibodies were diluted in 3% milk in TBS-X. To evaluate amyloid plaque pathology, tissue was incubated in HJ3.4 biotinylated antibody (anti-Aβ-1-13; 1.2μg/mL; a generous gift from the Holtzman lab). To assess NP-tau pathology, tissue was incubated in AT8 biotinylated antibody (anti-phospho-tau Ser202, Thr205; 1:500; ThermoFisher Scientific MN1020B). Sections were washed with TBS-X, then incubated in Vectastain ABC Elite solution (Vector Laboratories, PK-6100) in TBS for 1 hour at room temperature. Sections were washed with TBS and incubated in DAB solution. Sections were then mounted onto slides and cover slipped with Cytoseal XYL (Electron Microscopy Sciences, 18009). Brightfield imaging was performed with a Keyence BZ-X810 microscope.

All images were analyzed using NIH ImageJ software. For assessment of NP-tau pathology, the ipsilateral (representing NP-tau seeding) and contralateral (representing NP-tau spread) cortex and hippocampus were analyzed separately for each mouse. NP-tau seeding and spreading were expressed as a percentage of the total area for each brain region and averaged across three slices per animal. Statistical analyses were performed using GraphPad Prism 10. Differences in NP-tau seeding and spreading were measured using an unpaired Student’s t-test with Welch’s correction.

Plaque burden, plaque count, individual plaque size, and average plaque size were analyzed in the contralateral cortex and hippocampus for each mouse. Plaque pathology was expressed as a percentage of the total area for each brain region and averaged across two slices per animal. Plaque count was expressed as the number of plaques per mm^2^ averaged across two slices for each animal. Plaque size was expressed in µm^2^ averaged across two slices for each animal. Analysis of plaque distribution was performed by stratifying total plaque coverage based on size in µm^2^ and the frequency of occurrence in 349 um^2^ increments as previously described (Huynh, Liao et al. 2017). Statistical analyses were performed using GraphPad Prism 10. Differences in plaque burden, size, and count were measured using an unpaired Student’s t-test with Welch’s correction. Statistical difference in plaque distribution was measured using a two-way ANOVA.

### Immunofluorescence and X-34 plaque staining

Brain sections from each animal were washed three times with phosphate-buffered saline (PBS) then permeabilized with PBS containing 0.25% Triton-X100 for 30 minutes at room temperature. Sections were then stained with X-34 (10mM stock solution in DMSO) diluted 1:3000 in staining buffer containing 60% PBS and 40% ethanol, pH 10 for 20 minutes at room temperature. Sections were rinsed in a wash buffer containing 60% PBS and 40% ethanol. Following X-34 plaque staining, sections were blocked and permeabilized in PBS containing 0.4% Triton-X100 and 10% goat or donkey serum for 1 hour at room temperature, then incubated in primary antibodies diluted in PBS containing 0.4% Triton-X100 and 1% goat or donkey serum overnight at 4°C. The following primary antibodies were used: mouse anti-AT8b (ThermoFisher Scientific MN1020B; 1:500), goat anti-Iba1 (Abcam 5076; 1:100), rat anti-CD68 (BioRad MCA1957; 1:2000), rat anti-MHCII (BioLegend 107602; 1:50), rabbit anti-BACE1 (Abcam ab108394; 1:500), mouse anti-β-Amyloid 1-16 (AF647; 6E10; BioLegend 803021; 1:1000), and HJ3.4b anti-Aβ (a generous gift from the Holtzman lab; 1.2ug/mL). Sections were washed with PBS then incubated in secondary antibodies diluted 1:400 in PBS containing 0.25% Triton-X100 for 1 hour at room temperature. Secondary antibodies used in this study include goat anti-rabbit AF647 (Invitrogen A27040), donkey anti-rat AF488 (Invitrogen A21208), donkey anti-goat AF647 (Invitrogen A21447), and streptavidin AF647 (Invitrogen S21374). All secondary antibodies were diluted 1:400 in PBS containing 0.25% Triton-X100. Sections were washed with PBS, mounted on slides, and cover slipped with Fluoromount-G mounting medium (Invitrogen 00-4958-02).

### Confocal microscopy and image analysis

To assess peri-plaque NP-tau pathology (AT8), microglial recruitment (Iba1), microglial activation (CD68), microglial pro-inflammatory state (MHCII), neuritic dystrophy (BACE1), and plaque composition (6E10) in the cortex and hippocampus, a Zeiss LSM980 confocal microscope was used to acquire 20X z-stack images. Imaris was used to analyze images and make 3D reconstructions. NP-tau, Iba1, CD68, MHCII, and BACE1 measurements within 15μm of a plaque were classified as “peri-plaque” and summed per animal, then normalized to total X-34 volume per animal. To assess MHCII expression, MHCII fluorescence intensity was measured across full image volume and normalized to total image volume.

To assess plaque composition, the volumes of 6E10 and X-34 were calculated for each plaque. The X-34 proportion of the plaque volume was then calculated (X-34 volume/[X-34 volume + 6E10 volume – colocalized volume]).

Two sections per mouse were imaged with one image taken per brain region per section. All analyses were conducted in R (version 4.5.2). Outliers were identified and removed using the ROUT method (Q = 1%), and statistical comparisons were made in GraphPad Prism 10 using an unpaired Student’s t-test with Welch’s correction.

### Keyence imaging and analysis

To assess Iba1 intensity in the cortex and hippocampus, images were acquired using a Keyence BZ-X810 microscope. Laser intensity and exposure time were set by surveying tissue and selecting appropriate parameters that could remain the same for all samples. While these values varied by antibody, all sections in a staining cohort were imaged under identical conditions at the same magnification.

To determine Iba1 intensity, TIFF image files were opened in ImageJ and converted to 8-bit grayscale files. ROIs were drawn around the retrosplenial cortex and hippocampus, and the mean gray intensity of Iba1 was measured within each brain region. Measurements from two brain sections per mouse were analyzed and averaged for each brain region. Outliers were identified and removed using the ROUT method (Q = 1%), and statistical comparisons were made in GraphPad Prism 10 using an unpaired Student’s t-test with Welch’s correction.

## Results

### Ms4a4a loss remodels activated microglial states in the amyloid-bearing hippocampus

We previously demonstrated that *Ms4a4a* loss reduces amyloid pathology and modifies plaque-associated microglial phenotypes in 5xFAD mice at 6 months of age (Danhash, Verbeck et al. 2025). To determine how *Ms4a4a* loss alters microglial states and cellular programs within the amyloid-bearing hippocampus, we performed snRNA-seq on hippocampal tissue from non-transgenic (NTg), *Ms4a4a* KO (4A-KO), 5xFAD, and 5xFAD 4A-KO mice at 6 months of age (**Figure 1A**). Following quality control and integration, unsupervised clustering identified the major hippocampal cell populations, including excitatory neurons, inhibitory neurons, astrocytes, oligodendrocytes, oligodendrocyte precursor cells, endothelial cells, and microglia (**Figure 1B; Supplemental Table 2**). Comparison of overall cell-type proportions revealed that amyloid pathology was the dominant driver of cellular remodeling, with both 5xFAD groups exhibiting expansion of the microglial compartment relative to NTg and *Ms4a4a* KO mice, whereas loss of *Ms4a4a* alone had minimal effects on the overall cellular composition of the hippocampus (**Figure 1C**; Supplemental Table 2). To a lesser extent, we also observed a change in the astrocytes and vascular leptomeningeal cells (VLMCs) in both 5xFAD groups (**Figure 1C; Supplemental Table 2**).

**Figure 1.**
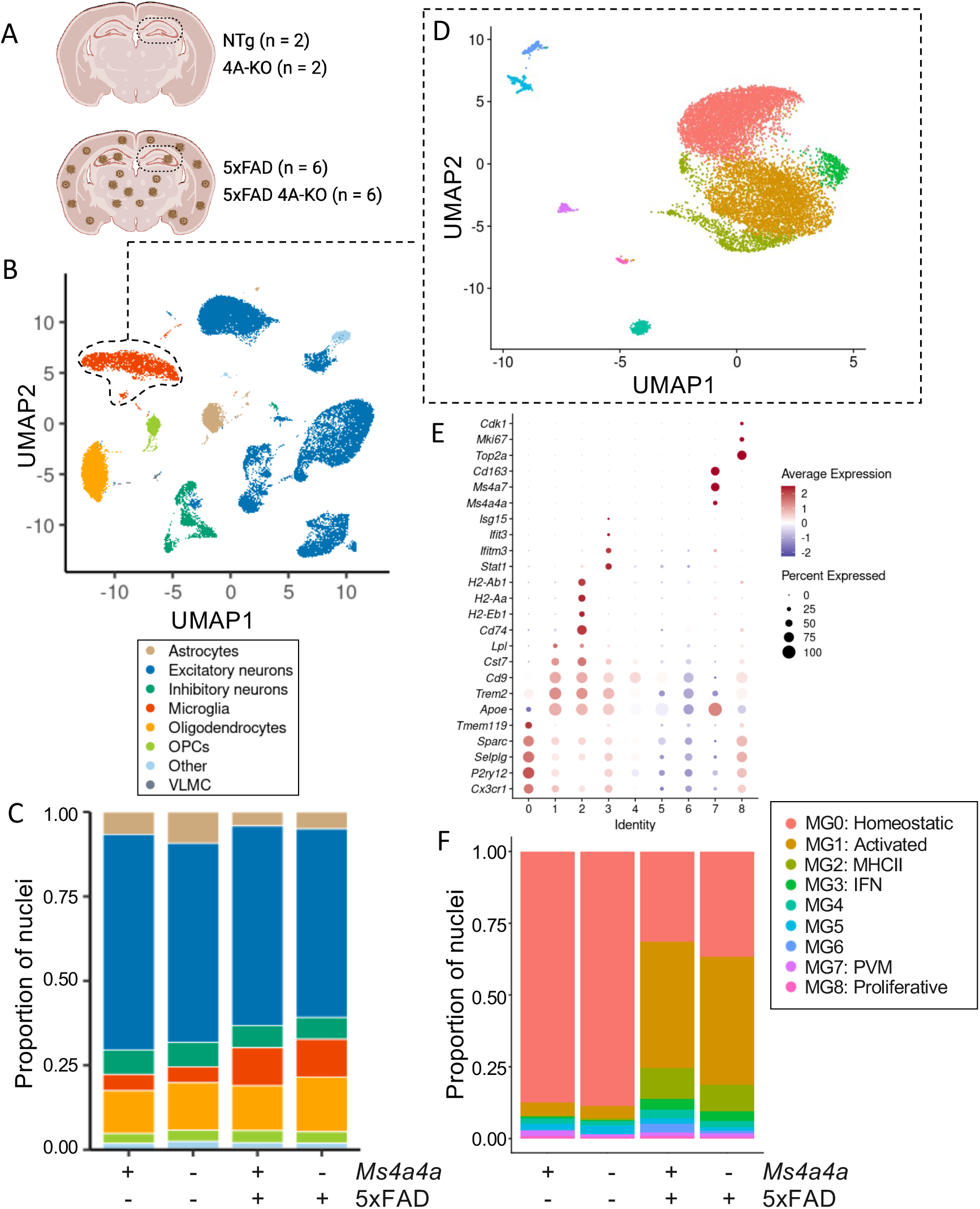
Ms4a4a loss does not broadly alter hippocampal cell composition but modifies disease-associated microglial programs *in vivo*. A. Study design. B. UMAP plot of nuclei isolated from hippocampi of 6-month-old 5xFAD and non-transgenic *Ms4a4a* WT and *Ms4a4a* KO mice (NTg, n = 2 males; *Ms4a4a* KO, n = 2 males; 5xFAD 4A-WT, n = 6 [50% female]; 5xFAD 4A-KO, n = 6 [50% female]). C. Cell-type proportions across genotypes. D. UMAP of microglia nuclei re-clustering which identified 9 distinct sub-types (MG0 – MG8). E. Subclusters were annotated based on gene expression of canonical markers. F. Proportion of nuclei contributing to each microglial subcluster across genotypes.

Given the prominent enrichment of *Ms4a4a* expression within microglia, we performed focused re-clustering of the microglial population. This analysis identified nine transcriptionally distinct microglial states defined by canonical marker genes associated with homeostatic, activated, interferon-responsive, HLAs, and proliferation, with *Ms4a4a* expression enriched in perivascular macrophages (PVM; **Figures 1D-E**). Across genotypes, the presence of amyloid pathology was the major determinant of microglial state composition (**Figure 1F; Supplemental Table 3**). Relative to NTg and *Ms4a4a* KO mice, 5xFAD animals exhibited a significant reduction in homeostatic microglia (MG0) accompanied by a significant expansion of activated (MG1), MHC-II (MG2), and interferon (MG3) microglial states (**Figure 1F; Supplemental Table 3**).

Ms4a4a loss in the setting of amyloid pathology did not substantially alter the overall distribution of major microglial states (**Figure 1F; Supplemental Table 3**), suggesting that Ms4a4a deletion does not broadly prevent the emergence of disease-associated microglia.

To determine whether *Ms4a4a* loss alters transcriptional programs within disease-associated microglial populations, we performed differential expression analyses comparing microglia from 5xFAD and 5xFAD 4A-KO mice within each identified microglial state (**Figure 2A**). Although *Ms4a4a* loss did not substantially alter the overall abundance of the major activated microglial states (Supplemental Table 3), widespread transcriptional changes were observed across multiple microglial subpopulations (**Figure 2B**), suggesting that *Ms4a4a* deletion reshapes microglial functional programs within the amyloid-bearing hippocampus rather than broadly preventing disease-associated state transitions.

**Figure 2:**
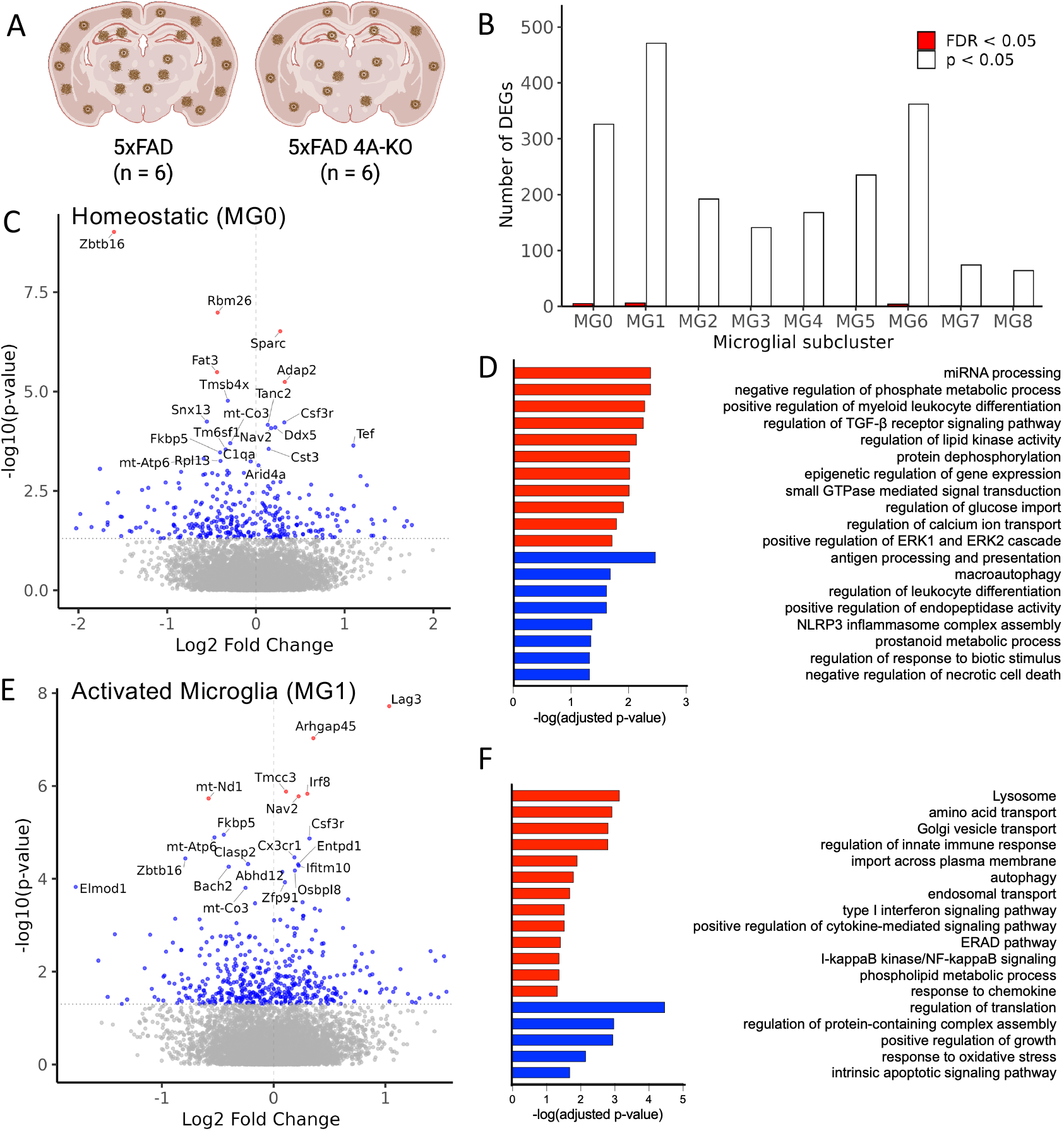
*Ms4a4a* loss alters transcriptional programs within homeostatic and activated microglia in amyloid-bearing mice. A. Schematic. B. Number of differentially expressed genes within major microglial subpopulations. Red, FDR<0.05. White, p<0.05. C-D. Homeostatic (MG0) cluster. C. Volcano plot of differentially expressed genes in the homeostatic microglia subcluster (MG0). Red, FDR < 0.05; blue, p < 0.05; gray, not significant. D. Pathway analysis of differentially expressed genes in homeostatic microglia (MG0; p<0.05). Pathways with an FDR<0.05 are shown. E-F. Activated (MG1) cluster. E. Volcano plot of differentially expressed genes in the activated microglia subcluster (MG1). Red, FDR < 0.05; blue, p < 0.05; gray, not significant. F. Pathway analysis of differentially expressed genes in activated microglia (MG1; p<0.05) reveals genes enriched in pathways related to immune signaling and proteostasis (FDR<0.05). Red, upregulated genes; blue, down-regulated genes.

Among the identified microglial states, the homeostatic (MG0) and activated (MG1) microglia clusters exhibited the largest number of differentially expressed genes following *Ms4a4a* loss (**Figure 2B**). Changes in the homeostatic (MG0) cluster suggested selective remodeling of the homeostatic microglial state in 5xFAD 4A-KO mice, including downregulation of *Zbtb16*, a negative regulator of pro-inflammatory activation, together with increased expression of *Sparc* and *Adap2*, genes associated with cellular morphology and cytoskeletal organization (**Figure 2C**; Supplemental Table 4). The homeostatic (MG0) cluster also exhibited differential expression of genes involved in metabolic regulation and transcriptional control (**Figure 2D**). We next examined transcriptional changes within the MG1 cluster, an activated microglial population enriched in amyloid-bearing mice. In 5xFAD 4A-KO microglia, genes associated with immune activation and enhanced cellular motility, including *Irf8*, *Arhgap45*, *Nav2*, and *Lag3*, were significantly upregulated relative to 5xFAD controls (**Figure 2E**; Supplemental Table 5). Pathway enrichment analyses of the activated (MG1) cluster revealed significant enrichment for immune signaling pathways, including chemokine signaling, type I interferon responses, and NF-κB signaling (**Figure 2F**). In parallel, multiple proteostatic and degradative pathways were also enriched, including autophagy, endosomal transport, lysosome organization, and endoplasmic reticulum-associated degradation, consistent with broad remodeling of microglial pathways involved in cellular stress responses and protein handling (**Figure 2E and 2F**). Importantly, prior studies implicate these pathways in the cellular handling and propagation of pathogenic protein aggregates (Cho, Cho et al. 2014, Heckmann, Teubner et al. 2019, Xu, Propson et al. 2021, Fleming, Bourdenx et al. 2022); thus, these findings raised the possibility that *Ms4a4a* loss may influence plaque-associated tau seeding and spreading within the amyloid-bearing brain. Together, these data demonstrate that while amyloid pathology is the dominant driver of microglial state transitions, *Ms4a4a* loss selectively remodels transcriptional states within activated microglial populations associated with inflammatory, lysosomal, and proteostatic response pathways.

### Ms4a4a loss selectively reduces dense-core, plaque-associated NP-tau accumulation

Having identified *Ms4a4a*-dependent remodeling in the microglial transcriptional programs associated with inflammatory and proteostatic pathways, we next sought to determine whether these transcriptional changes influence tau pathology in the amyloid-bearing brain. To do this, sarkosyl-insoluble tau aggregates isolated from human AD brain tissue (AD-tau) were stereotactically injected unilaterally into the dentate gyrus of the hippocampus and overlying cortex of 6-month-old 5xFAD and 5xFAD 4A-KO mice, and animals were aged for an additional three months prior to histological analysis at 9 months of age (**Figure 3A**).

**Figure 3.**
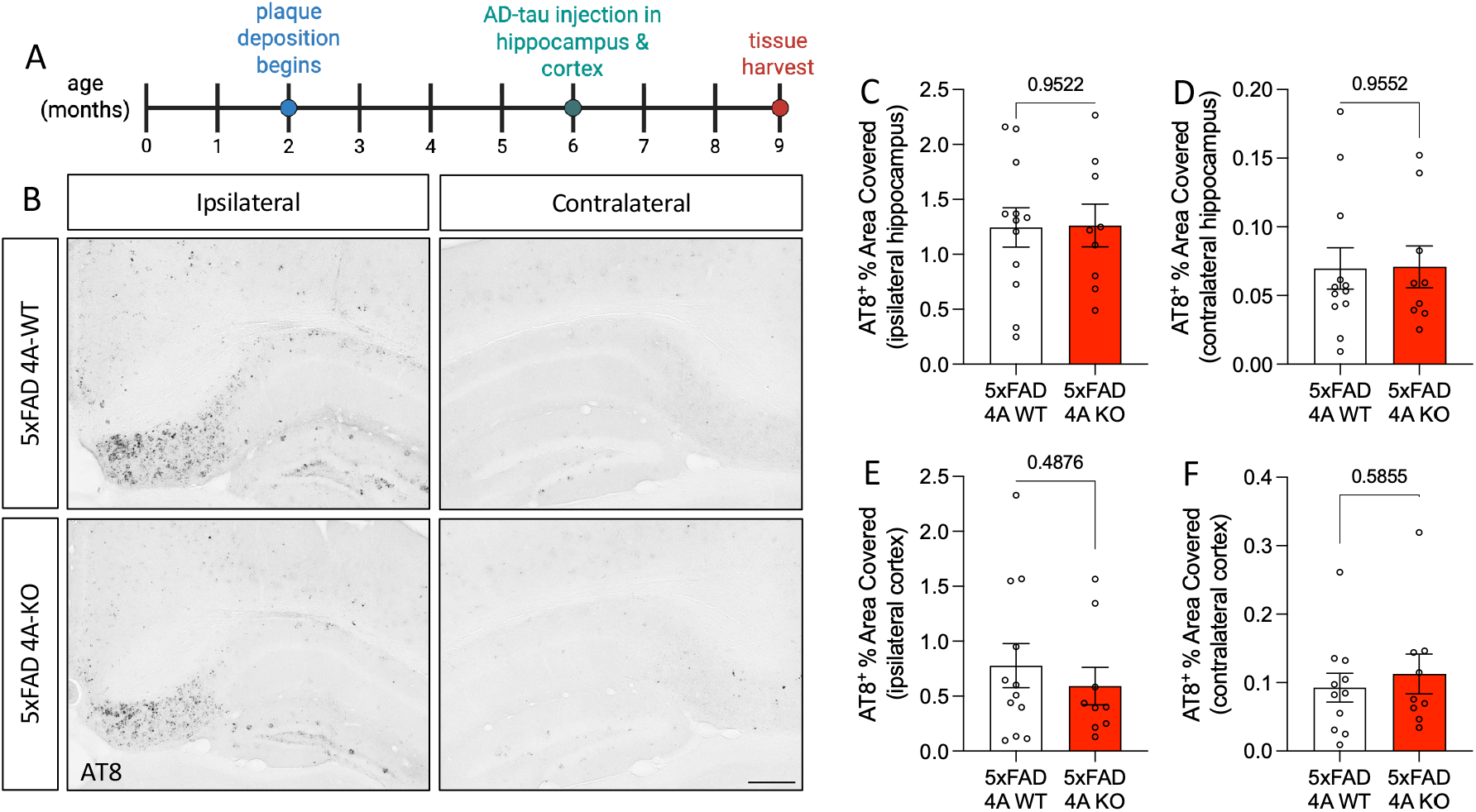
*Ms4a4a* loss does not alter overall phospho-tau pathology following AD-tau inoculation. A. Schematic depicting experimental timeline. B. Representative images of global neuritic plaque tau (NP-tau; AT8) in regions ipsilateral and contralateral to AD-tau injection of 5xFAD 4A-WT and 5xFAD 4A-KO mice. Scale bar, 250μm. C-D. Hippocampus. C. Quantification of global NP-tau in the ipsilateral hippocampus. D. Quantification of global NP-tau in the contralateral hippocampus. E-F. Cortex. E. Quantification of global NP-tau in the ipsilateral cortex. F. Quantification of global NP-tau in the contralateral cortex. Data represent 5xFAD 4A-WT n = 12; 5xFAD 4A-KO n = 9; female mice. Each dot represents one animal (mean of 3 brain sections). Graphs represent mean ± standard error of the mean (SEM). Statistical comparisons were made using an unpaired Student’s t-test with Welch’s correction. Outliers identified and removed using the ROUT method (Q = 1%; F).

We first quantified overall phosphorylated tau pathology using AT8 immunostaining across the hippocampus and cortex. Robust AT8-positive pathology was detected in both 5xFAD and 5xFAD 4A-KO mice following AD-tau inoculation (**Figure 3B**). Quantification of total AT8 immunoreactivity revealed no significant differences between genotypes in either the ipsilateral or contralateral hippocampus or cortex (**Figure 3C-F**), indicating that loss of *Ms4a4a* does not broadly suppress global phospho-tau seeding, spreading, or accumulation following tau inoculation.

To determine whether *Ms4a4a* loss selectively alters plaque-associated tau pathology, we quantified NP-tau surrounding amyloid deposits following AD-tau inoculation. NP-tau is defined based on its localization near amyloid plaques (He, Guo et al. 2018, Leyns, Gratuze et al. 2019, Gratuze, Chen et al. 2021, Chen, Song et al. 2024); so, we first analyzed plaque-associated AT8 pathology using X-34 labeling to identify dense-core fibrillar plaques (defined as AT8-positive volume within 15μm of an X-34-positive fibrillar plaque core; **Figure 4A**). In the ipsilateral cortex, corresponding to the primary site of tau injection and local seeding, 5xFAD 4A-KO mice exhibited a trend toward less plaque-associated NP-tau relative to 5xFAD controls (p=0.09; **Figure 4A-B**). In contrast, analysis of the contralateral cortex, a region reflecting tau spreading, propagation, and accumulation of tau pathology, demonstrated significantly less in X- 34-associated NP-tau in 5xFAD 4A-KO mice (p=0.01; **Figure 4A and 4C**). NP-tau near fibrillar plaques did not differ between groups in the hippocampus (Supplemental Figure 1A-C). Thus, we conclude that *Ms4a4a* loss reduces the accumulation of fibrillar plaque-associated NP-tau at sites distal to initial tau inoculation.

**Figure 4.**
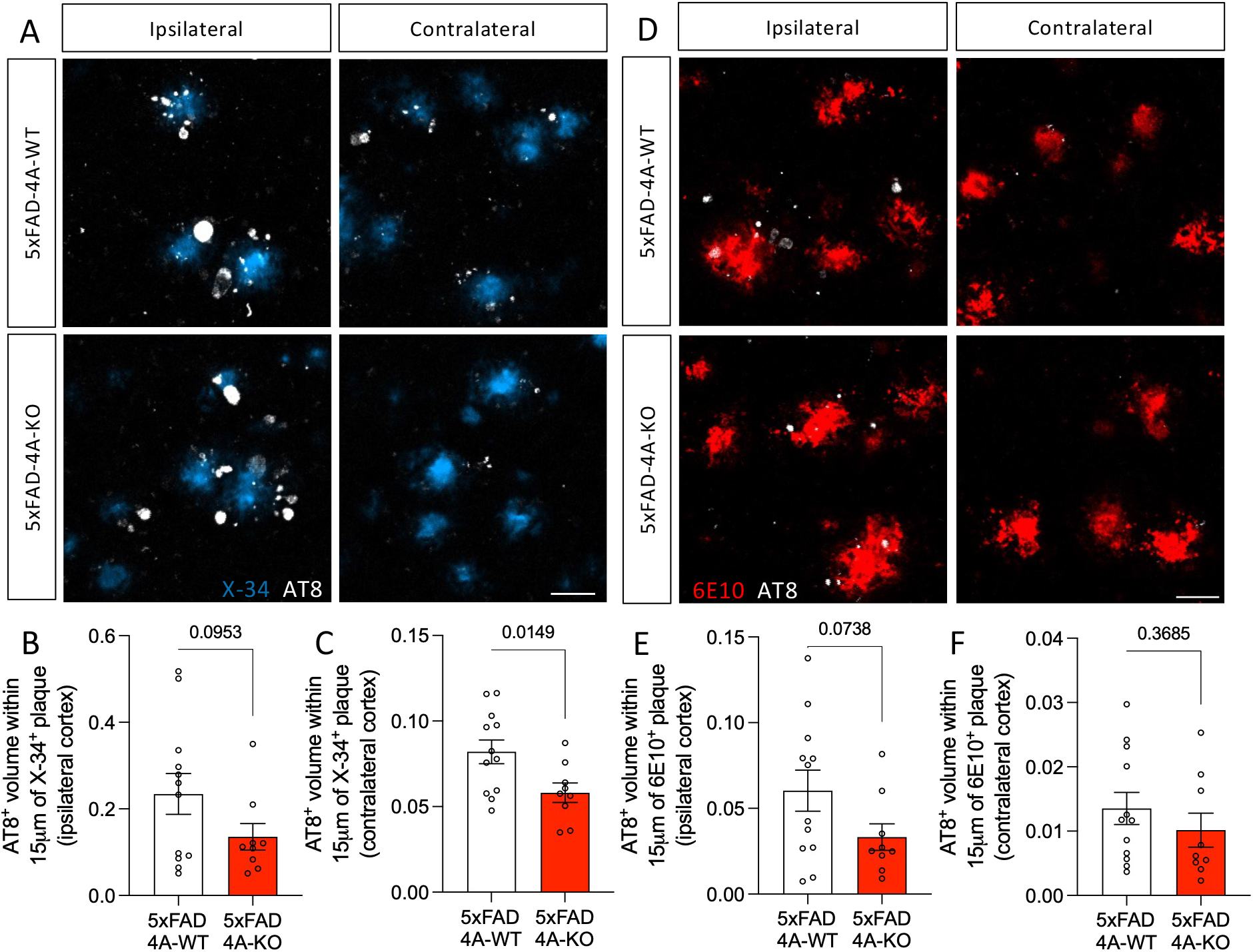
*Ms4a4a* loss selectively reduces dense-core plaque-associated NP-tau accumulation. A-C. Evaluation of NP-tau near fibrillar amyloid plaques labeled with X-34. A. Representative immunofluorescence confocal images of fibrillar Aβ plaque core (X-34; blue) and peri-plaque NP-tau (AT8; white) in the cortex in regions ipsilateral and contralateral to NP-tau injection. Scale bar, 20μm. B. Quantification of peri-plaque (fibrillar) NP-tau in ipsilateral cortex. C. Quantification of peri-plaque (fibrillar) NP-tau in the contralateral cortex. D-F. Evaluation of NP-tau near total amyloid plaques labeled with 6E10. D. Representative immunofluorescence confocal images of total Aβ plaques (6E10; red) and peri-plaque NP-tau (AT8; white) in the ipsilateral and contralateral cortex. Scale bar, 20μm. E. Quantification of peri-plaque (total) NP-tau in ipsilateral cortex. F. Quantification of peri-plaque (total) NP-tau in contralateral cortex. Data represent 5xFAD 4A-WT n = 12; 5xFAD 4A-KO n = 9; female mice. Each dot represents one animal (mean of 2 brain sections). Graphs represent mean ± SEM. Statistical comparisons were made using an unpaired Student’s t-test with Welch’s correction.

We next sought to determine whether the reduction in plaque-associated NP-tau extended across amyloid plaque populations. Total amyloid plaques were evaluated using 6E10 immunolabeling (**Figure 4D**). Similar to the X-34 analysis, quantification of 6E10-associated NP-tau revealed a trend toward reduced ipsilateral plaque-associated AT8 pathology in 5xFAD 4A-KO mice in the cortex (p=0.07; **Figure 4E**). However, in contrast to the X-34-defined, dense-core plaque analysis, contralateral 6E10-associated NP-tau was not significantly different between genotypes (p=0.36; **Figure 4F**). NP-tau near 6E10 was unchanged in the hippocampus (Supplemental Figure 1D-F). The distinct associations between global AT8 burden and plaque-associated NP-tau across both plaque populations is consistent with the spatial restriction of the effect, as dense-core plaque-proximal NP-tau represents a subpopulation of total phospho-tau pathology, and reductions confined to the plaque microenvironment may be insufficient to produce a detectable change in global AT8 immunoreactivity. Together, these findings demonstrate that *Ms4a4a* loss does not broadly reduce overall phospho-tau pathology, but instead selectively attenuates NP-tau accumulation within certain dense-core plaque-associated microenvironments. The stronger effect observed using X-34-defined fibrillar plaques suggests that *Ms4a4a*-dependent microglial programs may preferentially influence tau pathology associated with mature dense-core amyloid deposits.

### Ms4a4a loss does not substantially alter plaque-associated microglial activation following AD-tau inoculation

*Ms4a4a* expression is enriched in microglia (Novikova, Kapoor et al. 2021, Silva-Gomes, Mapelli et al. 2022, Brase, You et al. 2023), which are key brain phagocytes involved in the uptake, processing, and clearance of extracellular tau and contribute to tau pathology (Bolos, Llorens-Martin et al. 2016, Gratuze, Chen et al. 2021). Microglia also form a protective barrier around Aβ plaques, mitigating neuritic dystrophy where AD-tau seeds amplify (Condello, Yuan et al. 2015, He, Guo et al. 2018, Leyns, Gratuze et al. 2019, Sadleir, Gomez et al. 2025). To determine whether the lesser burden of dense-core plaque-associated NP-tau associated with *Ms4a4a* loss was accompanied by altered plaque-associated microglial responses, we quantified microglial abundance and activation surrounding amyloid plaques following AD-tau inoculation. Brain sections were co-labeled with X-34 to identify dense-core fibrillar plaques together with Iba1 and CD68 to assess plaque-associated microglia and activation, respectively (Figure 5A; Supplemental Figure 2A). Quantification of plaque-associated Iba1 immunoreactivity revealed no significant difference between 5xFAD and 5xFAD 4A-KO mice in the ipsilateral and contralateral cortex (**Figure 5B–C**) and hippocampus (Supplemental Figure 2B–C). Thus, *Ms4a4a* loss does not substantially alter the overall recruitment or accumulation of microglia surrounding dense-core plaques following AD-tau inoculation. Similarly, CD68 immunoreactivity surrounding X-34-positive plaques was comparable between genotypes in both hemispheres (**Figure 5D–E; Supplemental Figure 2D-E**), suggesting that the reduction in plaque-associated NP-tau was not accompanied by major changes in plaque-associated microglial activation at the histologic level.

**Figure 5.**
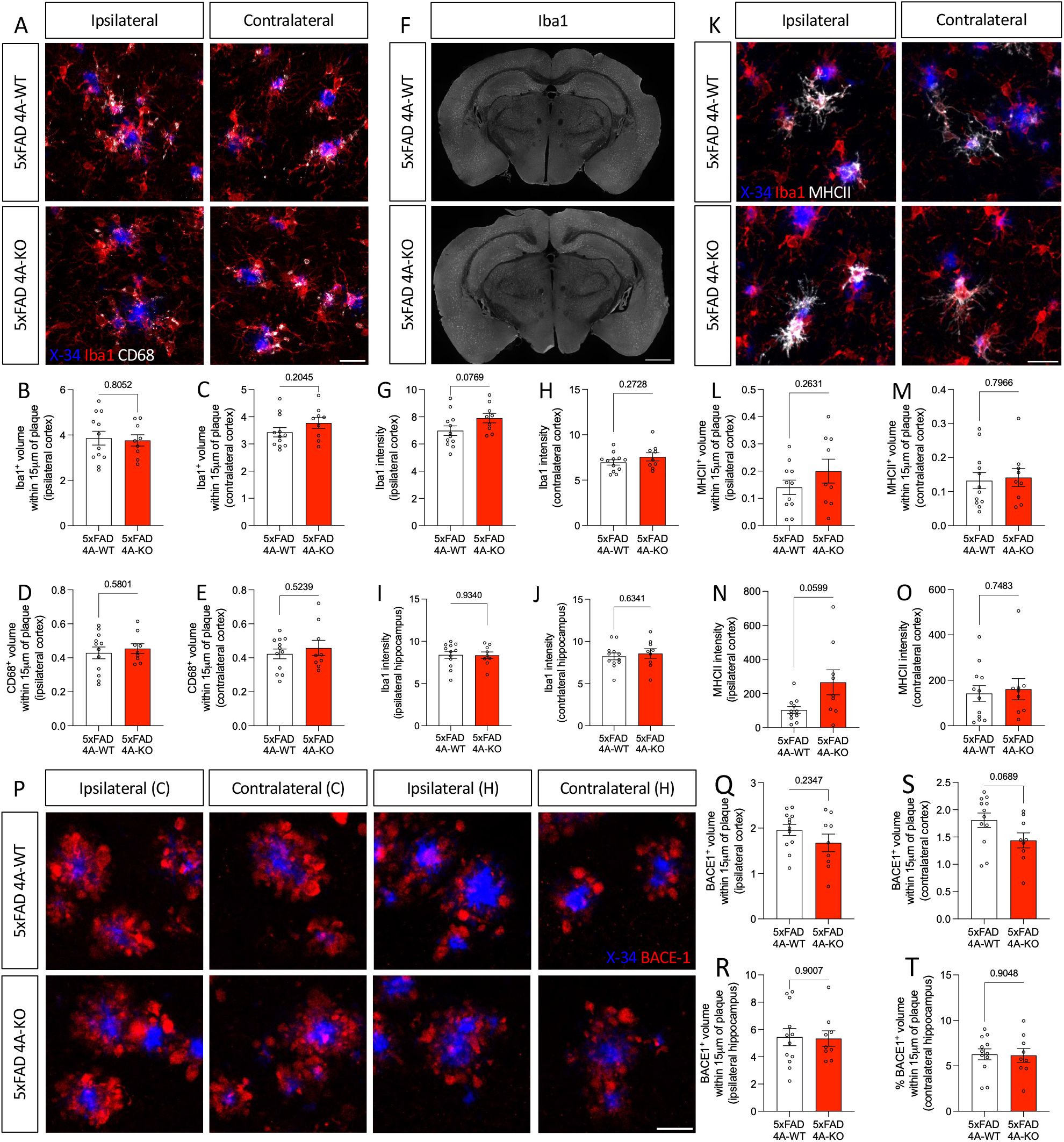
Reduced dense-core plaque-associated NP-tau occurs without major changes in plaque-associated microglial accumulation or activation with *Ms4a4a* loss. A. Representative immunofluorescence confocal images of fibrillar Aβ plaques (X-34; blue), total microglia (Iba1; red), and a reactive microglia marker (CD68; white) in the ipsilateral and contralateral cortex. Scale bar, 20μm. B. Quantification of Iba1-positive microglial volume within 15μm of plaques in the ipsilateral cortex. C. Quantification of Iba1-positive microglial volume within 15μm of plaques in the contralateral cortex. D. Quantification of CD68-positive volume within 15μm of plaques in the ipsilateral cortex. E. Quantification of CD68-positive volume within 15μm of plaques in the contralateral cortex. F. Representative images of microglia (Iba1; white) in whole tissue sections. Scale bar, 1000μm. G. Quantification of intensity of Iba1 in the ipsilateral cortex. H. Quantification of intensity of Iba1 in the contralateral cortex. I. Quantification of intensity of Iba1 in the ipsilateral hippocampus. J. Quantification of intensity of Iba1 in the contralateral hippocampus. K. Representative immunofluorescence confocal images of fibrillar Aβ plaques (X-34; blue), total microglia (Iba1; red), and a proinflammatory microglia marker (MHCII; white) in the ipsilateral and contralateral cortex. Scale bar, 20μm. L. Quantification of MHCII-positive volume within 15μm of plaques in the ipsilateral cortex. M. Quantification of MHCII-positive volume within 15μm of plaques in the contralateral cortex. N. Quantification of MHCII intensity normalized to image volume in the ipsilateral cortex. O. Quantification of MHCII intensity normalized to image volume in the contralateral cortex. P-T. Ms4a4a loss modestly reduces neuritic dystrophy. P. Representative immunofluorescence confocal images of fibrillar Aβ plaques (X-34; blue) and dystrophic neurites (BACE1; red) in the ipsilateral and contralateral cortex and hippocampus. Scale bar, 20μm. C, cortex. H, hippocampus. Q. Quantification of BACE1-positive volume within 15μm of plaques in the ipsilateral cortex. R. Quantification of BACE1-positive volume within 15μm of plaques in the ipsilateral cortex. S. Quantification of BACE1-positive volume within 15μm of plaques in the contralateral hippocampus. T. Quantification of BACE1-positive volume within 15μm of plaques in the contralateral hippocampus. Data represent 5xFAD 4A-WT n = 12; 5xFAD 4A-KO n = 9; female mice. Each dot represents one animal (mean of 2 brain sections). Graphs represent mean ± SEM. Statistical comparisons were made using an unpaired Student’s t-test with Welch’s correction. Outliers identified and removed using the ROUT method (Q = 1%; L, N).

Because snRNA-seq analyses identified prominent transcriptional remodeling of activated microglial states following *Ms4a4a* loss, we next assessed whether broader regional microglial activation was altered after AD-tau inoculation. Quantification of total Iba1 intensity, a proxy for microglial activation, demonstrated a trend toward increased overall Iba1 expression within the ipsilateral cortex of 5xFAD 4A-KO mice relative to 5xFAD controls (p=0.07; **Figure 5F–G**). Total Iba1 intensity in the contralateral cortex (**Figure 5H**) and hippocampus (**Figure 5I-J**) were unchanged.

Activation and recruitment to plaques represent only a subset of microglial function. Our group previously observed that *Ms4a4a* loss shifts microglia to a pro-inflammatory state in the context of amyloid pathology (Danhash, Verbeck et al. 2025). To determine whether *Ms4a4a* deficiency alters the inflammatory state of microglia in the context of AD-tau inoculation, brain sections from 5xFAD 4A-WT and 5xFAD 4A-KO mice injected with AD-tau were stained for Aβ plaques (X-34), total microglia (Iba1), and pro-inflammatory microglia (MHCII; **Figure 5K**, cortex; Supplemental Figure 2F, hippocampus). Expression of the antigen presentation marker, MHCII, surrounding amyloid plaques remained unchanged between genotypes (**Figure 5K–M**; Supplemental Figure 2F-H), further indicating that *Ms4a4a* loss does not broadly enhance classical plaque-associated inflammatory activation following tau injection. Yet, the overall MHCII expression was significantly elevated in the ipsilateral cortex (p=0.05; **Figure 5N**) and unchanged in the contralateral cortex (**Figure 5O**) and hippocampus (Supplemental Figure 2I-J). Together, these findings demonstrate that reduced dense-core plaque-associated NP-tau accumulation in 5xFAD 4A-KO mice occurs in the absence of major changes in plaque-associated microglial abundance, CD68 enrichment, or overt plaque-associated MHCII expression. Instead, these data suggest that *Ms4a4a* loss more selectively alters microglial functional states and proteostatic programs rather than inducing large-scale changes in plaque-associated microglial recruitment or overt inflammatory activation.

Aβ plaques cause local neuritic dystrophy, with smaller plaques resulting in proportionately more neuritic dystrophy (Condello, Yuan et al. 2015, Sadleir, Popovic et al. 2022). As peri-plaque dystrophic neurites provide the ideal niche for NP-tau seeding and later spreading (He, Guo et al. 2018), we sought to assess whether *Ms4a4a* loss impacted plaque-associated neuritic dystrophy in 5xFAD mice using BACE1 immunolabeling. Quantification of BACE1-positive dystrophic neurites surrounding X34-positive plaques revealed no significant differences in the ipsilateral cortex or hippocampus between 5xFAD and 5xFAD 4A-KO mice (**Figure 5P–R**). In the contralateral cortex, 5xFAD 4A-KO mice exhibited a modest reduction in plaque-associated BACE1 immunoreactivity that did not reach statistical significance (p = 0.06; **Figure 5S**), while contralateral hippocampal BACE1 levels were unchanged (**Figure 5T**). Together, these findings suggest that reduced dense-core plaque-associated NP-tau associated with *Ms4a4a* loss occurs in the absence of major reductions in plaque-associated dystrophic neurite pathology.

### Ms4a4a loss does not substantially alter amyloid plaque burden or composition following AD-tau inoculation

We previously reported that loss of *Ms4a4a* reduces amyloid plaque burden and alters plaque-associated microglial responses in 6-month-old 5xFAD mice (Danhash, Verbeck et al. 2025). To determine whether the reduction in dense-core plaque-associated NP-tau observed in 5xFAD 4A-KO mice at 9 months of age was accompanied by persistent alterations in amyloid pathology, we next quantified amyloid plaque burden and composition following AD-tau inoculation. Brain sections were analyzed using complementary amyloid labeling approaches to assess total plaque burden, plaque morphology, and fibrillar plaque composition within the hippocampus and cortex (**Figure 6**; Supplemental Figure 3).

**Figure 6.**
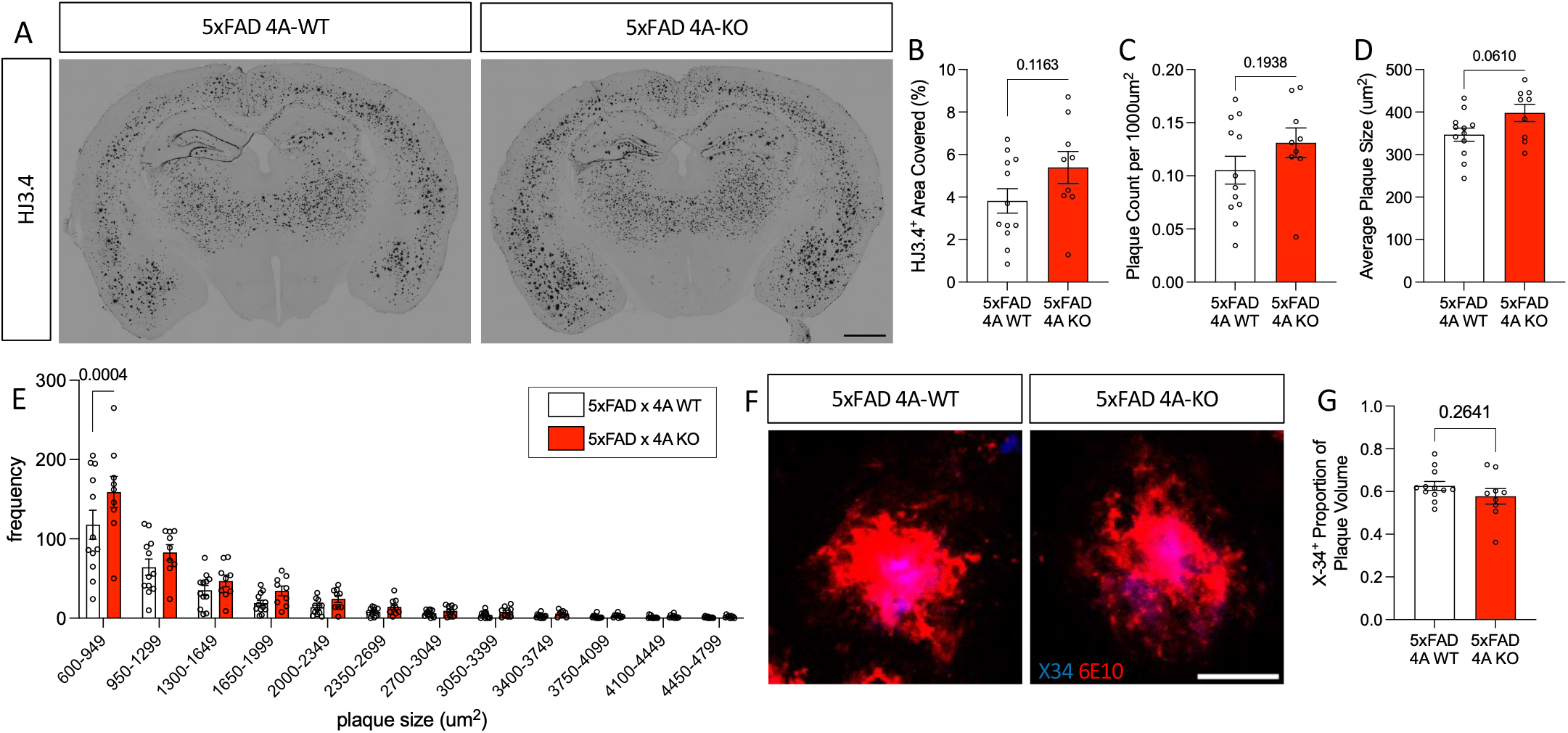
Ms4a4a loss alters plaque morphology without substantially changing late-stage amyloid burden or fibrillar plaque composition. A. Representative images of immunohistochemistry for Aβ plaques (HJ3.4). Scale bar, 1000μm. B. Quantification of HJ3.4 percent area within the cortex. C. Quantification of average number of plaques/1000μm^2^ within the cortex. D. Quantification of average plaque size within the cortex (μm^2^). E. Quantification of the frequency of plaques binned based on size in μm^2^ within the cortex. F. Representative immunofluorescence confocal images of fibrillar Aβ plaque core (X-34; blue) and total Aβ (6E10; white) in the cortex. Scale bar, 20μm. G. Quantification of X-34 proportion relative to total plaque volume. Data represent 5xFAD 4A-WT n = 12; 5xFAD 4A-KO n = 9; female mice. Each dot represents one animal (mean of 2 brain sections). Graphs represent mean ± SEM. Statistical comparisons were made using an unpaired Student’s t-test with Welch’s correction (B-D, G) and two-way ANOVA (E).

In contrast to our prior findings at 6 months of age, overall amyloid plaque burden was not significantly altered in 9-month-old 5xFAD 4A-KO mice following AD-tau inoculation. Quantification of total amyloid plaque area revealed no significant differences between 5xFAD and 5xFAD 4A-KO mice (**Figure 6A-B; Supplemental Figure 3A**), and total plaque number was similarly unchanged between genotypes (Figure 6C; Supplemental Figure 3B). Analysis of plaque morphology, however, demonstrated region-specific alterations in plaque size distribution. Average plaque size was modestly increased in the cortex of 5xFAD 4A-KO mice relative to 5xFAD controls (p = 0.06; **Figure 6D**) and significantly increased in the hippocampus (p = 0.03; Supplemental Figure 3C). In parallel, size distribution analyses revealed a significantly greater abundance of smaller plaques (600–949 μm²) in both the cortex (p = 0.0004; **Figure 6E**) and hippocampus (p = 0.0012; Supplemental Figure 3D) of 5xFAD 4A-KO mice.

Because the reduction in NP-tau was most pronounced surrounding X-34-positive dense-core plaques, we next assessed whether *Ms4a4a* loss altered amyloid plaque composition. We previously reported that 5xFAD 4A-KO mice exhibit increased plaque compaction at 6 months of age (Danhash, Verbeck et al. 2025). In contrast, quantification of fibrillar versus total amyloid plaque labeling at 9 months revealed no significant differences in plaque composition between genotypes in the cortex (**Figure 6F-G**) and a significant decrease in plaque compaction in the hippocampus (p = 0.02; Supplemental Figure 3E-F), suggesting that *Ms4a4a* loss alters the relative abundance of dense-core amyloid deposits at this later disease stage in a region-specific manner.

Together, these findings demonstrate that, in contrast to the reduced amyloid burden and altered plaque compaction previously observed in 6-month-old 5xFAD 4A-KO mice, *Ms4a4a* loss does not substantially alter overall amyloid plaque burden, plaque number, or fibrillar plaque composition at 9 months following AD-tau inoculation. Instead, *Ms4a4a* loss was associated with more subtle alterations in plaque morphology, including increased average plaque size and enrichment of smaller plaques. Importantly, the reduction in dense-core plaque-associated NP-tau observed in 5xFAD 4A-KO mice occurred in the absence of major changes in bulk amyloid deposition. These findings suggest that *Ms4a4a* loss may selectively alter plaque-associated microenvironmental or microglial processes linked to NP-tau accumulation rather than broadly suppressing amyloid deposition during later stages of disease progression.

## Discussion

In this study, we demonstrated that loss of *Ms4a4a* selectively remodels activated microglial programs and alters cortical dense-core plaque-associated tau pathology in the amyloid-bearing brain. Genetic variants within the *MS4A* locus are strongly associated with AD risk and resilience and have increasingly been linked to microglial biology (Deming, Filipello et al. 2019, Novikova, Kapoor et al. 2021, Wightman, Jansen et al. 2021, Brase, You et al. 2023, You, Brase et al. 2023, Wang, Nykanen et al. 2024, Danhash, Verbeck et al. 2025, Rosner, Sun et al. 2026), although their functional consequences for plaque-associated tau pathology remain poorly understood (Hansen, Karch et al. 2026). We found that amyloid pathology remained the dominant driver of microglial state transitions; yet, *Ms4a4a* deficiency in the setting of amyloid pathology shifted activated microglial transcriptional programs toward interferon, lysosomal, and proteostatic pathways. In parallel, *Ms4a4a* loss reduced cortical dense-core plaque-associated NP-tau spread following AD-derived tau inoculation despite relatively modest changes in overall phospho-tau burden, plaque-associated microgliosis, and late-stage amyloid burden. These findings suggest that *Ms4a4a*-dependent microglial states selectively influence plaque-associated microenvironments linked to local tau pathology rather than broadly suppressing amyloid or tau accumulation. Our study supports a model in which AD resilience-associated microglial pathways shape downstream plaque-associated neurodegenerative processes during later stages of amyloid pathology.

Microglial activation is a prominent feature of AD and is increasingly recognized as a major contributor to disease heterogeneity and progression (Hamelin, Lagarde et al. 2016, Hansen, Hanson et al. 2018, Sun, Victor et al. 2023). While the presence of amyloid pathology was the dominant driver of microglial state transitions in our study, loss of *Ms4a4a* selectively remodeled gene expression programs within activated microglial populations rather than broadly altering microglial state proportions. Activated microglia from 5xFAD 4A-KO mice exhibited enrichment of pathways linked to interferon signaling, lysosomal organization, autophagy, endosomal transport, and ER-associated degradation, supporting a role for *Ms4a4a* in regulating proteostatic and inflammatory microglial programs in the setting of amyloid pathology. These pathways are consistent with disease-associated microglial programs linked to cellular stress responses, antigen presentation, and protein handling pathways observed across amyloid models and human AD brain tissue (Chen, Lu et al. 2020, Roy, Chiu et al. 2022, Choi, Wang et al. 2023, Sun, Victor et al. 2023, Mancuso, Fattorelli et al. 2024). These findings are consistent with emerging studies implicating MS4A family proteins in microglial effector functions associated with TREM2 signaling, lysosomal biology, and disease-associated microglial states (Deming, Filipello et al. 2019, Wang, Fan et al. 2022, Rosner, Sun et al. 2026). Interestingly, although snRNA-seq analyses pointed to substantial transcriptional remodeling, histologic analyses revealed relatively modest changes in plaque-associated Iba1, CD68, and MHCII immunoreactivity. Together, these findings suggest that *Ms4a4a* loss primarily alters microglial functional states and proteostatic pathways rather than inducing large-scale changes in plaque-associated microglial recruitment or overt inflammatory activation. This dissociation between transcriptional remodeling and overt histologic microgliosis further supports the concept that microglial functional states may influence disease progression through selective modulation of local plaque-associated microenvironments.

Here, we show that *Ms4a4a* loss selectively reduced cortical dense-core plaque-associated NP-tau accumulation following AD-tau inoculation. Importantly, this phenotype was not accompanied by reductions in overall phospho-tau burden, suggesting that *Ms4a4a*-dependent microglial programs preferentially influence localized plaque-associated tau pathology rather than broadly suppressing tau accumulation throughout the hippocampus. The reduction in NP-tau was strongest when analyses were restricted to X34-positive dense-core fibrillar plaques and was less pronounced when all amyloid deposits were assessed using 6E10 labeling. These findings suggest that mature fibrillar plaque microenvironments may differentially regulate plaque-associated tau accumulation and neuritic injury. Compact fibrillar plaque cores are associated with plaque-associated neuritic dystrophy and NP-tau accumulation, suggesting that plaque architecture may influence the formation of local seeding-competent microenvironments (He, Guo et al. 2018, Sadleir, Popovic et al. 2022, Tsering, Hery et al. 2023, Sadleir, Gomez et al. 2025). Thus, age- and region-dependent changes in plaque morphology following *Ms4a4a* loss may alter the availability of plaque-associated niches that support local tau accumulation. Dense-core plaques are closely associated with dystrophic neurites, activated microglia, and local proteostatic stress, supporting a model in which plaque architecture and plaque-associated cellular responses shape the local accumulation of phosphorylated tau (He, Guo et al. 2018, Clayton, Delpech et al. 2021, Tsering, Hery et al. 2023). Interestingly, the most robust reduction in NP-tau was observed in the contralateral hemisphere distal to the initial tau inoculation site, consistent with altered processes linked to plaque-associated tau propagation or secondary accumulation. However, because global AT8 burden remained unchanged, our findings more strongly support selective modulation of plaque-associated tau microenvironments rather than broad suppression of tau seeding. Although plaque-associated BACE1-positive dystrophic neurites were largely preserved following *Ms4a4a* loss, we observed a modest reduction in contralateral cortical dystrophic neurites that paralleled the reduction in dense-core plaque-associated NP-tau. These findings suggest that selective modulation of local neuritic injury may contribute to altered plaque-associated tau pathology independently of major changes in plaque burden. However, the AD-tau inoculation paradigm models amyloid-associated tau seeding and propagation rather than the full spectrum of primary tauopathy. It will therefore be important to determine whether *Ms4a4a*-dependent microglial programs similarly influence tau pathology in models lacking amyloid deposition.

These findings also extend our prior work demonstrating that *Ms4a4a* deficiency alters amyloid pathology in 5xFAD mice. Previously, we reported that 6-month-old 5xFAD 4A-KO mice exhibit reduced amyloid plaque burden together with increased plaque compaction (Danhash, Verbeck et al. 2025). In contrast, at the later 9-month timepoint examined following AD-tau inoculation, we observed minimal changes in overall amyloid burden, plaque number, or fibrillar plaque composition. Instead, *Ms4a4a* loss was associated with more subtle alterations in plaque morphology, including increased average plaque size and enrichment of smaller X34-positive plaques. Notably, the reduction in dense-core plaque-associated NP-tau persisted despite the absence of major late-stage reductions in bulk amyloid pathology. These findings suggest that the impact of *Ms4a4a* loss evolves across disease progression, with earlier effects on amyloid accumulation giving way to more selective modulation of plaque-associated microenvironments during later stages of pathology, consistent with stage-dependent effects reported for other microglia-enriched AD risk genes such as *TREM2* (Meilandt, Ngu et al. 2020). More broadly, these data support the concept that plaque-associated tau pathology may become partially dissociated from bulk amyloid burden as disease progresses (Hojjati, Chiang et al. 2024). These findings are consistent with studies demonstrating that microglial ablation or disruption of microglia-enriched AD risk pathways alters amyloid-associated tau seeding and propagation (Leyns, Gratuze et al. 2019, Gratuze, Chen et al. 2021, Wang, Fan et al. 2022, Chen, Song et al. 2024). Together, these observations support a model in which microglial functional states actively shape the influence of amyloid pathology on downstream tau accumulation rather than simply reflecting ongoing disease processes.

Several observations from this study provide insight into potential mechanisms linking *Ms4a4a*-dependent microglial programs to NP-tau accumulation. Activated microglial states from 5xFAD 4A-KO mice were enriched for pathways associated with lysosomal organization, autophagy, endosomal transport, and ER-associated degradation, suggesting that altered proteostatic responses may influence local plaque-associated tau accumulation. Among genes altered within activated microglia, *Lag3* was increased in association with *Ms4a4a* loss. *LAG3* has been implicated in the uptake and propagation of aggregation-prone proteins, including α-synuclein and tau, although the contribution of microglial *LAG3*-associated pathways to plaque-associated tau pathology remains poorly understood (Chen, Kumbhar et al. 2024, Yang, Jeong et al. 2025, Zhang, You et al. 2025, Yang, Kumbhar et al. 2026). Because *Lag3* induction occurred alongside enrichment of interferon and proteostatic pathways, these findings may reflect broader remodeling of activated plaque-associated microglial states rather than a direct effect of *LAG3* itself on NP-tau accumulation. The distinct pathological effects observed following *Ms4a4a* loss compared to *Trem2* deficiency may further reflect the positioning of MS4A4A upstream of TREM2-associated microglial signaling pathways (Deming, Filipello et al. 2019, Wang, Nykanen et al. 2024, Rosner, Sun et al. 2026). Although plaque-associated microgliosis and dystrophic neurite pathology were largely preserved, it remains possible that *Ms4a4a* loss alters more subtle aspects of microglial-neuritic interactions, local proteostasis, or plaque-associated extracellular environments that were not captured by static histologic analyses. Because these studies utilized global *Ms4a4a* knockout mice, we cannot exclude contributions from peripheral myeloid populations to the observed effects on plaque-associated tau pathology. Future studies incorporating microglia-specific manipulations, longitudinal analyses, and functional interrogation of proteostatic pathways will be important for defining how *Ms4a4a*-dependent microglial states influence plaque-associated tau pathology.

## Conclusions

Together, our findings identify *Ms4a4a* as a regulator of plaque-associated microglial programs linked to NP-tau accumulation in the amyloid-bearing brain. More broadly, this work supports a model in which AD resilience-associated microglial pathways selectively shape plaque-associated microenvironments that promote downstream tau pathology. These findings have important therapeutic implications, suggesting that modulation of microglial functional states may alter pathogenic plaque-associated tau processes even in the absence of major reductions in bulk amyloid burden. Because common AD-associated variants occur within or near the *MS4A* locus, determining how MS4A4A regulates plaque-associated microglial states may help clarify how AD risk and resilience variants influence downstream neurodegenerative processes. Defining the mechanisms through which microglial pathways regulate local plaque-associated proteostatic environments may reveal therapeutic strategies aimed at selectively limiting downstream neurodegenerative processes in AD.

## Acknowledgments

We thank Alector for providing the *Ms4a4a* KO mice used in this study. We thank Dalya Rosner, Torri Ball, Timothy Miller, and Miwei Hu for thoughtful discussions. This work was supported by access to equipment made possible by the Hope Center for Neurological Disorders, the Neurogenomics and Informatics Center, and the Departments of Neurology and Psychiatry at Washington University School of Medicine. Diagrams were generated using BioRender.com.

## Consent Statement

Not applicable.

## Authors’ contributions

Designed experiments: EPD, CMK. Performed and analyzed experiments: EPD, SYF, JAM, RDA, ACV, GH, SFY, WKS, CMK. Provided reagents: RJP, WKS, DMH. Provided funding: CMK. Wrote the manuscript: EPD, CMK. Revised and approved manuscript: EPD, SYF, JAM, RDA, ACV, GH, SFY, EEF, RJP, WKS, DMH, CMK.

