## Supplemental Figure 1 for "*Ms4a4a* loss reprograms amyloid-associated microglia and limits dense-core plaque-associated tau spreading"

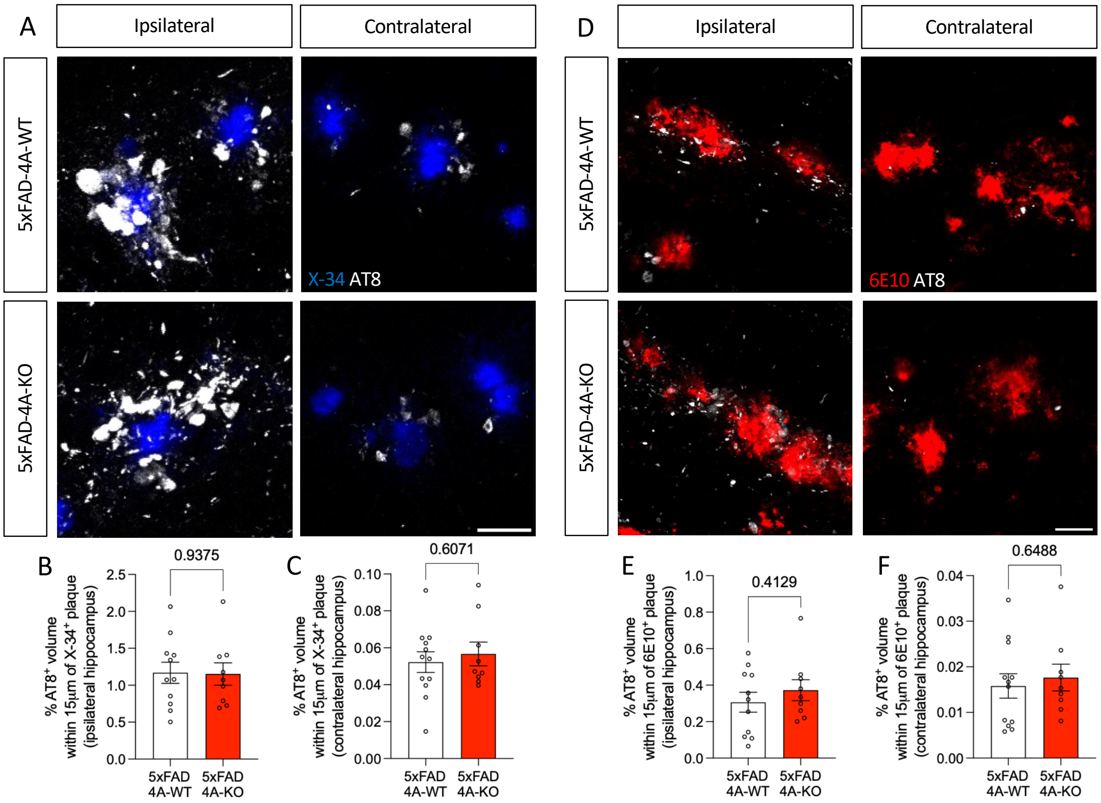


**Supplemental Figure 1. Loss of *Ms4a4a* does not alter local NP-tau pathology in the hippocampus.** A. Representative immunofluorescence confocal images of fibrillar Aβ plaque core (X-34; blue) and peri-plaque NP-tau (AT8; white) in the ipsilateral and contralateral hippocampus. Scale bar, 20μm. B. Quantification of peri-plaque NP-tau in ipsilateral hippocampus. C. Quantification of peri-plaque NP-tau in contralateral hippocampus. D. Representative immunofluorescence confocal images of total Aβ plaques (6E10; red) and peri-plaque (total) NP-tau (AT8; white) in the ipsilateral and contralateral hippocampus. Scale bar, 20μm. E. Quantification of peri-plaque (total) NP-tau in ipsilateral hippocampus. F. Quantification of peri-plaque (total) NP-tau in contralateral hippocampus. Data represent 5xFAD 4A-WT n = 12; 5xFAD 4A-KO n = 9; female mice. Each dot represents one animal (mean of 2 brain sections). Graphs represent mean ± SEM. Statistical comparisons were made using an unpaired Student's t-test with Welch's correction. Outliers identified and removed using the ROUT method (Q = 1%; B, E).
