## Supplemental Figure 2 for "*Ms4a4a* loss reprograms amyloid-associated microglia and limits dense-core plaque-associated tau spreading"

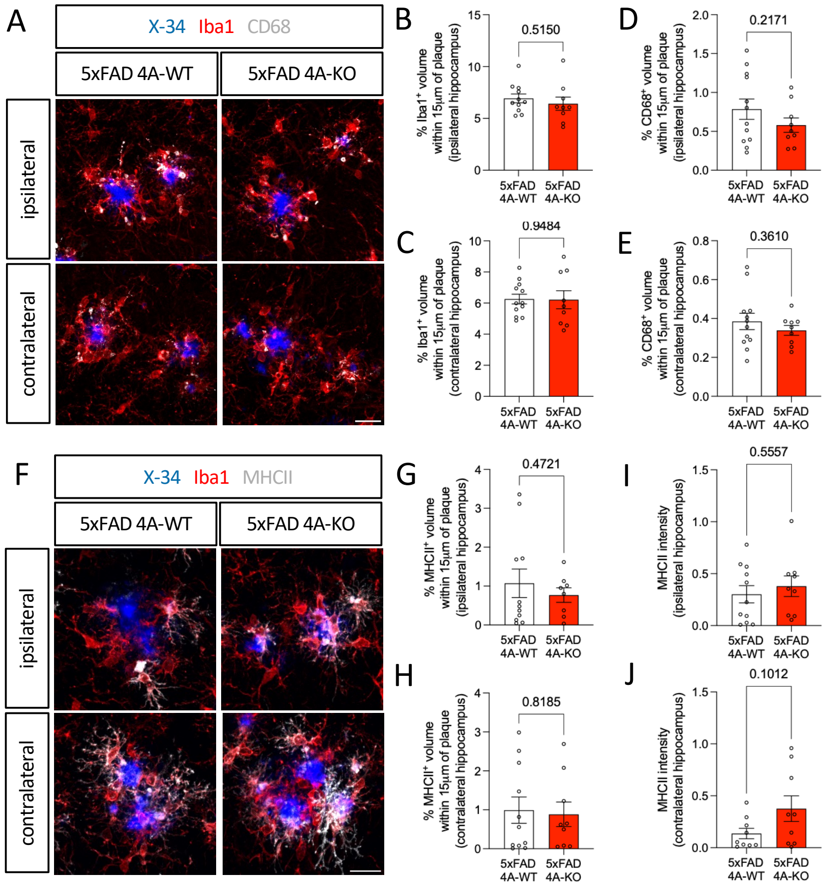


**Supplemental Figure 2. Loss of *Ms4a4a* does not alter microglial recruitment or reactivity in the hippocampus.** A. Representative immunofluorescence confocal images of fibrillar Aβ plaques (X-34; blue), total microglia (Iba1; red), and a reactive microglia marker (CD68; white) in the ipsilateral and contralateral hippocampus. Scale bar, 20μm. B. Quantification of Iba1-positive microglial volume within 15μm of plaques in the ipsilateral hippocampus. C. Quantification of Iba1-positive microglial volume within 15μm of plaques in the contralateral hippocampus. D. Quantification of CD68-positive volume within 15μm of plaques in the ipsilateral hippocampus. E. Quantification of CD68-positive volume within 15μm of plaques in the contralateral hippocampus. F. Representative immunofluorescence confocal images of fibrillar Aβ plaques (X-34; blue), total microglia (Iba1; red), and a proinflammatory microglia marker (MHCII; white) in the ipsilateral and contralateral hippocampus. Scale bar, 20μm. G. Quantification of MHCII-positive volume within 15μm of plaques in the ipsilateral hippocampus. H. Quantification of MHCII-positive volume within 15μm of plaques in the contralateral hippocampus. I. Quantification of MHCII intensity normalized to microglia volume in the ipsilateral hippocampus. J. Quantification of MHCII intensity normalized to microglia volume in the contralateral hippocampus. Data represent 5xFAD 4A-WT n = 12; 5xFAD 4A-KO n = 9; female mice. Each dot represents one animal (mean of 2 brain sections). Graphs represent mean ± SEM. Statistical comparisons were made using an unpaired Student's t-test with Welch's correction. Outliers identified and removed using the ROUT method (Q = 1%; B, G-J).
