## Supplemental Figure 3 for "*Ms4a4a* loss reprograms amyloid-associated microglia and limits dense-core plaque-associated tau spreading"

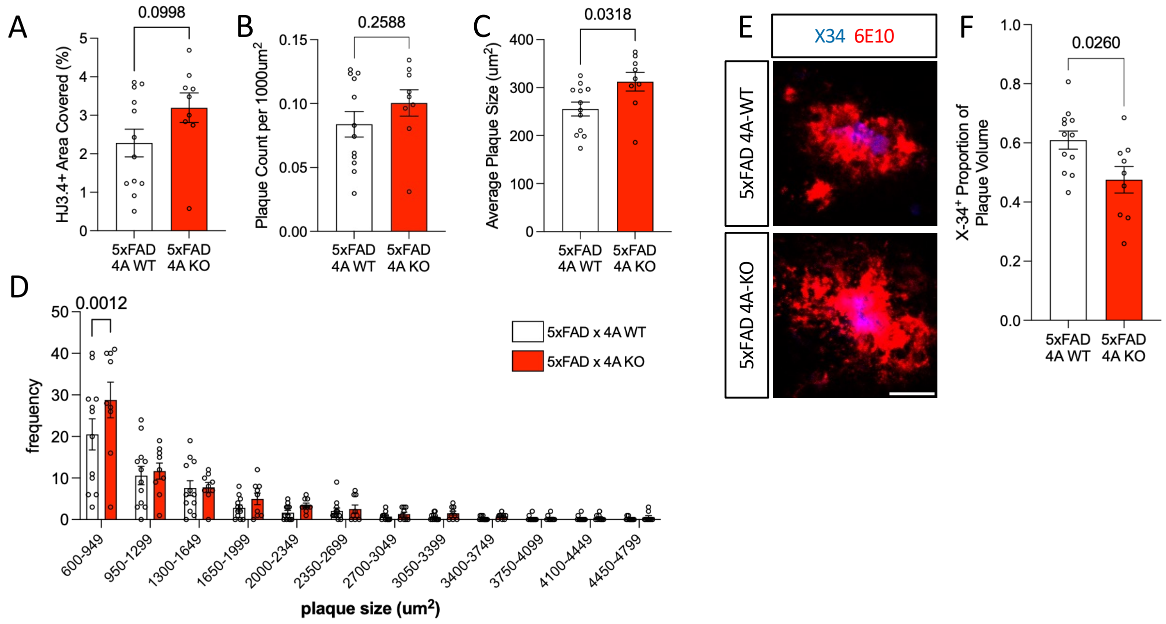


**Supplemental Figure 3.** ***Ms4a4a loss alters plaque morphology without substantially changing late-stage amyloid burden or fibrillar plaque composition in the hippocampus*****.** A. Quantification of HJ3.4 percent area within the hippocampus. B. Quantification of average number of plaques/1000μm$^2$ within the hippocampus. C. Quantification of average plaque size within the hippocampus (μm$^2$). D. Quantification of the frequency of plaques binned based on size in μm$^2$ within the hippocampus. E. Representative immunofluorescence confocal images of fibrillar Aβ plaque core (X-34; blue) and total Aβ (6E10; white) in the hippocampus. Scale bar, 20μm. F. Quantification of X-34 proportion relative to total plaque volume in the hippocampus. Data represent 5xFAD 4A-WT n = 12; 5xFAD 4A-KO n = 9; female mice. Each dot represents one animal (mean of 2 brain sections). Graphs represent mean ± SEM. Statistical comparisons were made using an unpaired Student's t-test with Welch's correction (A-C, F) and two-way ANOVA (D).
